# Impact of prenatal synthetic glucocorticoid exposure on the adolescent brain

**DOI:** 10.1101/2023.04.14.536872

**Authors:** Ricardo Magalhães, Nuno Gonçalves, Rui Sousa, Ana Coelho, Carina Soares-Cunha, Pedro Moreira, Paulo Marques, Jetro J Tuulari, Nora M Scheinin, Linnea Karlsson, Hasse Karlsson, Nuno Sousa, Ana João Rodrigues

**Author notes:** Corresponding authors: Ana João Rodrigues, PhD, & Nuno Sousa, MD PhD, University of Minho, ICVS | School of Medicine, Campus de Gualtar, 4710-057 Braga, Portugal. Current affiliation: NeuroSpin, CEA, CNRS, Paris-Saclay University, Gif-sur-Yvette, France. Current affiliation: Departamento de Psiquiatria e Saúde Mental, Centro Hospitalar Tondela-Viseu, Viseu, Portugal (Rui Sousa). Contributed equally to this work (Ricardo Magalhães, Nuno Gonçalves, Rui Sousa).

## Abstract

Synthetic Glucocorticoids (sGC) are commonly prescribed in preterm risk pregnancies in order to improve fetal organ maturation. This administration greatly reduces perinatal and neonatal mortality and respiratory distress syndrome associated with prematurity, but preclinical evidence warns for an adverse effect of sGC in the developing brain.

In this work we evaluated the long-term effects of prenatal exposure to sGC in the brain of 17 years-old adolescents using multimodal MRI. From 4607 birth registrations from Hospital de Braga - Portugal, we selected participants that were born with similar gestational age, but that were either exposed during pregnancy to sGC (n=21) or non-exposed (n=24). After obtaining a detailed clinical history, participants were subjected to an extensive neuropsychological evaluation, followed by structural and functional MRI.

No differences were found in the performance on neuropsychological tests between sGC-exposed and non-exposed participants. Moreover, no differences were found in regional brain volumes. However, the sGC-exposed group presented reduced functional connectivity at rest in a network involving primarily sub-cortical, cerebellar and frontal nodes in comparison to the non-exposed group, even after controlling for confounding factors such as gestational age at birth, birth weight, and sex.

Our results suggest that prenatal sGC-exposed adolescents present no significant deviations in neuropsychological performance in the dimensions that we evaluated, although they presented altered functional connectivity, highlighting the need for additional studies to understand the impact of these changes in brain functioning and in behavior.

**Highlights:** Prenatal synthetic glucocorticoid exposure does not lead to structural changes in the adolescent brain.

Adolescents prenatally exposed to synthetic glucocorticoids present altered resting state network.

## 1. Introduction

Maternal stress hormones, glucocorticoids (GC), are critical to normal fetal development, increasing in concentration over pregnancy to ensure fetal maturation and to prepare for birth (Mastorakos and Ilias, 2003). Since GC are potent hormones with pleiotropic physiological effects, they are tightly regulated by the placental enzyme 11β-hydroxysteroid dehydrogenase type 2 (11β-HSD2) that degrades GC into inactive metabolites (Benediktsson et al., 1997; Seckl and Holmes, 2007). This control is crucial for normative fetal development, as increased levels of these hormones can increase the risk for adverse health outcomes later in life.

Synthetic glucocorticoids (sGC) are often administered to pregnant women at risk of premature labour to promote fetal lung maturation and to prevent respiratory distress syndrome in preterm infants. Prenatal administration of sGC is known to increase surfactant production and alveolar volume and accelerates the maturation of other organs, such as the kidney, liver and heart, reducing most of the severe outcomes of prematurity. Current guidelines indicate that prenatal sGC therapy is adequate for women at risk of premature delivery before 34 weeks of gestation. The actual recommended treatment includes one single course of sGC, consisting of either two 12mg doses of betamethasone 24 hours apart or four 6mg doses of dexamethasone every 12 hours. However, these dosages are empirical, as no systematic studies define the minimum dose to induce the same beneficial outcomes. Betamethasone and dexamethasone are used interchangeably and are commonly preferred because they are not readily metabolized by placental 11β-HSD2 and are 25 times more potent than endogenous GC (Ballard and Ballard, 1995; Brown et al., 1996; Chapman et al., 2013).

Although it is clear that sGC administration has decreased neonatal mortality and morbidity (McGoldrick et al., 2020), increasing evidence pinpoints potential deleterious effects of these hormones in the developing brain. Animal studies have shown that prenatal sGC exposure induces structural and functional changes in different brain regions such as the prefrontal cortex, hippocampus, nucleus accumbens, and amygdala, amongst others (Caetano et al., 2017; Cartier et al., 2016; Duarte et al., 2019; McArthur et al., 2005; Oliveira et al., 2006, 2011, 2012; Reynolds, 2013; Rodrigues et al., 2012; Soares-Cunha et al., 2014). These changes can be the result of a direct effect of sGC on the developing brain, but also a consequence of an impaired hypothalamic-pituitary-adrenal (HPA) axis, often observed in models of prenatal stress or GC exposure (Braun et al., 2013; Moisiadis and Matthews, 2014a, 2014b; Reynolds, 2013). These structural and functional changes are often associated with the behavioral deficits observed in animal models of sGC exposure. For example, we have shown that prenatal dexamethasone exposure in late gestation alters the structure and function of nucleus accumbens neurons, and that optogenetic or pharmacological modulation of these neurons rescues the motivational deficits observed in this model (Soares-Cunha et al., 2014, 2016).

In humans the literature about the impact of prenatal sGC is sparse and controversial. Different studies have suggested that prenatal sGC treatment alters the HPA-axis, leading to altered stress reactivity later in life (Braun et al., 2013; Moisiadis and Matthews, 2014a, 2014b; Reynolds, 2013). HPA-axis de-regulation can partially underlie the reported structural and functional alterations, as it has been shown that sGC can program neuronal migration and maturation in non-human primates (Coe and Lubach, 2005). Studies with multiple courses of prenatal sGC treatment (previous administration guidelines) showed an association with lower birth weight and decreased head circumference, decreased cortical volume and complexity of cortical folding (Asztalos et al., 2014; Crowther et al., 2019; Modi et al., 2001; Murphy et al., 2001, 2008; Wapner et al., 2006). These findings suggest that prenatal sGC can indeed lead to enduring structural changes. Importantly, multiple courses of prenatal sGC did not improve pre-term birth outcomes, and were associated with increased risk for neurodevelopmental and neuropsychiatric conditions later in life (Asztalos et al., 2014; French et al., 2004; Melamed et al., 2019; Murphy et al., 2008; Räikkönen et al., 2020) providing evidence against the use of multiple courses. Still, it is important to refer that other reports present no evidence of altered neurodevelopment (Crowther et al., 2007; Wapner et al., 2006).

Even fewer studies evaluate the potential effects of the current recommended single course of sGC on brain structure and function. One study has shown that administration of betamethasone was associated with acute changes in higher cortical functions in the exposed fetuses using magnetoencephalography (Schneider et al., 2011). In another study, children (6-10 years) with fetal sGC exposure had prominent cortical thinning in the right anterior cingulate cortex (ACC) (Davis et al., 2013). Because a thinner right ACC is associated with risk for affective problems, the authors postulated that this phenotype could arise later in life. In agreement, others have shown that the prevalence of any mental or behavioral disorders was higher in sGC-exposed groups independently of prematurity (Räikkönen et al., 2020; Wolford et al., 2019). Conversely, other studies reported no association between prenatal sGC and worse child neurodevelopmental outcomes (Alexander et al., 2016; Davis et al., 2013; Stutchfield et al., 2013). The contradictory findings are likely the result of different study designs, differences in outcome definitions, and measurements as well as differences in the timing of sGC administration as well as associated maternal or fetal conditions.

It is clear that human studies on the effects of prenatal sGC exposure on brain development are very limited, highlighting the need to perform additional studies to understand the long-term impact of these hormones in the fetus. In this study, we analyzed a cohort of healthy 17-year-old adolescents that were prenatally exposed to sGC and non-exposed individuals. We performed: i) a detailed clinical characterization; ii) neuropsychological evaluation, focusing on emotionality; and iii) structural, functional and diffusion MRI acquisition.

## 2. Materials and Methods

### 2.1. Ethics Statement

The study here described was conducted following the principles stated in the Declaration of Helsinki and was approved by the ethics committee of Hospital de Braga and the School of Medicine of Universidade do Minho. To participate in the study, all participants and legal tutors gave written informed consent after being explained the aims and procedures of the study. Participants were informed that they could withdraw from the study at any time.

### 2.2. Study design

The target population were adolescents that were prenatally exposed to one single course of sGCs and with no history of serious health conditions and no history of familiar neuropsychiatric disorders. A non-exposed group was recruited based on individuals with similar age, sex, gestational weight and maternal age. The recruitment was performed by trained researchers at the Clinical Academic Centre (CCA) – Braga (https://www.ccabraga.org), Portugal.

Recruitment had different phases: phase one – inspection of the physical birth records of Hospital de Braga and posterior selection of potential participants based on exclusion and inclusion criteria; phase 2 – participants’ contact by telephone to explain the rationale of the project and invitation to participate; phase 3 – visit to the recruitment center CCA, located at Hospital de Braga, and beginning of the study. After detailed explanation of the study, informed consent was signed by one of the parents/legal tutors and the adolescent. The study was divided into two sessions. Session 1) details of the medical records were confirmed with the parents, and additional relevant events were included. Maternal and paternal sociodemographic and pertinent clinical information was also obtained. The mother answered a questionnaire regarding gestation, birth and the neonatal and childhood periods to obtain additional information that was not previously registered in the Hospital/Health records. In this session we also included a detailed neuropsychological evaluation (see below). Session 2) Adolescents that were eligible for the study were invited to undergo multimodal MRI.

As a compensation for their participation in the study, all adolescents (even the ones excluded in session 1) were offered the opportunity to visit the laboratories of ICVS/School of Medicine University of Minho and interact with neuroscientists during this period.

### 2.3. Participant details

The study population consists of individuals born in Hospital de Braga (Braga, Portugal) during the years of 2000 and 2001. Of 4774 birth records, only 76.41% had information that could be validated physically (paper records). The other clinical registrations were either missing, incomplete or transferred to other health units. We double checked clinical records of mothers and children included in the cohort to cross-validate information, especially with regard to glucocorticoid administration at any moment in life (phase 1).

Collected maternal information consisted of sociodemographic information, maternal age at birth, delivery method, unifetal vs gemelar gestation, number of previous pregnancies, number of previous births, pregnancy intercurrences (if any), relevant clinical history and history of medication. In the cases of mothers that were administered with sGC, the following data was also obtained: GCs administration day, dosage and number of administrations and reason for administration.

Fetal information included: sex, gestational age, weight and height at birth, neonatal intercurrences (if any) and medication, especially sGC administration in the neonatal period.

Inclusion criteria: exposure to a single course of sGC in the prenatal period (GC-exposed) and sex-matched control individuals (non-exposed).

Exclusion criteria: major fetal structural abnormalities; chorioamnionitis; chronic maternal condition or medication during pregnancy (including GCs administration of any type); sGC administration in the post-natal period; use of steroids for other aims; use of maternal glucocorticoids during lactation; preterm with <34 weeks of gestation. We also excluded all participants with familial or individual history of psychiatric disorders.

### 2.4. Psychological Measures

For the neuropsychological-psychiatric evaluation, the study population consisted of 21 individuals within the GCs group and 24 within the control group, forming a total of 45 subjects.

The following questionnaires were used:

- NEO-FFI (Bertoquini and Pais-Ribeiro, 2006) to measure of the five domains of personality - Neuroticism, Extraversion, Openness, Agreeableness, and Conscientiousness;
- Beck depression inventory (BDI-II) (Beck et al., 1996) to evaluate depressive symptomatology;
- Beck anxiety inventory (BAI) (Beck et al., 1988) to evaluate anxious traits;
- Perceived Stress Scale 10 (PSS10) (Cohen et al., 1983) for individual stress levels;
- Strengths and Difficulties Questionnaire (SDQ) (Goodman, 1997) for a short behavioral screening (Emotional symptoms, Conduct problems, Hyperactivity/inattention, Peer relationship problems, Prosocial behavior)
- Substance Use Risk Profile Scale (SURPS) (Canfield et al., 2015) by assessing four well-validated personality risk factors for substance misuse - Impulsivity, Sensation Seeking, Anxiety Sensitivity, and Hopelessness;
- Behavioral Inhibition Scale/Behavioral Avoidance Scale (BIS/BAS), that measures two motivational systems: motivation to avoid aversive outcomes or to approach goal-oriented outcomes (Carver and White, 1994).

Previously coded questionnaires were fulfilled by the participant alone, placed in a closed envelope and sent for analysis by another researcher.

### 2.5. Statistical analysis

Data was analyzed using SPSS (version 24.0;IBM). First, we assessed the normality of distribution of the data. Comparisons of means between groups were analyzed by parametric independent-samples t-test (for normal distribution) or non-parametric Mann-Whitney U-tests.

Details for MRI data analysis is provided in the next section.

### 2.6. MRI acquisitions

For the MRI study, several subjects were excluded for the following reasons: orthodontic appliances; extensive movement or artifacts during the acquisition; adolescent decided to quit during the acquisition. Finally, 17 individuals within the GCs group and 11 individuals within the control group were included, resulting in a total of 28 subjects.

All imaging sessions were performed in Hospital de Braga on a clinically approved Siemens Magnetom Avanto 1.5T MRI Scanner (Siemens, Erlagen, Germany). A 12-channel receive only head coil was used. The scanning protocol included three sequences: as structural acquisition, a T1 weighted Magnetization Prepared Rapid Gradient Echo (MPRAGE) sequence with TR=2730 ms, TE=3,48 ms, 176 sagittal slices, 1 mm^3^ isometric resolution over a 256×256 matrix; a resting-state functional MRI (rs-fMRI) acquisition using an Echo Planar Imaging (EPI) sequence with TR=2000 ms, TE= 30 ms, slice thickness = 4 mm, slice gap = 0.48mm, in plane voxel size = 3.5 x 3.5 mm^2^, over a 65 x 65 matrix, with 30 axial slices and 190 volumes; and a diffusion weighted imaging (DWI) acquisition using a spin-echo echo-planar imaging (SE-EPI) sequence with the following parameters: TR=8800 ms, TE=99 ms, FoV=240×240 mm, acquisition matrix=120×120, 61 2-mm axial slices with no gap, 30 non-collinear gradient direction with b=1000 s/mm-2, one b=0 s/mm-2 and a total of 2 repetitions. During the rs-fMRI acquisition, all participants were instructed to remain still with eyes closed and letting their mind wonder freely without focusing on anything in specific.

A trained neuro-radiologist visually inspected all acquisitions ensuring none presented any lesions or pathologies requiring exclusion from the study. Subjects were also excluded when they presented movement exceeding the voxel size or critical artifacts (Soares et al., 2016).

### 2.7. MRI data-processing and analysis

Processing of structural data for volumetric analysis was performed using the Freesurfer pipeline (v5.10, surfer.nmr.mgh.harvard.edu), which has been already described in detail (Desikan et al., 2006; Fischl et al., 2002). The process performs 31 processing steps, including registration to Talairach space, skull stripping, intensity normalization, tessellation of different tissue classes and segmentation of sub-cortical, cortical and white matter areas. The pipeline has been validated against manual segmentations (Fischl et al., 2002). Based on the literature, and for the current analysis, 7 sub-cortical regions of the aseg atlas (Fischl et al., 2002) were chosen from each hemisphere, namely the Thalamus, Caudate, Putamen, Pallidum, Hippocampus, Amygdala and the Accumbens, as well as 5 regions from each hemisphere of the Destrieux cortical parcellation based on the functional results, including the anterior and posterior cingulum, superior frontal cortex, cerebellar gray matter and the precuneus, leading to a total of 24 regions of interest included in the analysis.

Preprocessing of rs-fMRI data was executed using FMRIB Software Library (FSL, v6.0, https://fsl.fmrib.ox.ac.uk/fsl/fslwiki). The preprocessing steps included: slice-timing correction using the costume Siemens interleaved order; motion correction; registering all volumes to the mean image; motion outlier detection (using *fslmotion_outliers* with default metrics and parameters and subjects were excluded if over 15% of the volumes were detected as outliers); brain extraction to create individual brain masks for the functional and structural data; calculation of the normalization parameters from the native functional space to the standardized MNI space in three steps by 1) calculating the linear registration parameters from the native functional space to the native structural space, 2) calculating the linear and the non-linear registration parameters from the native structural space to the standardized MNI space, 3) combining the calculated parameters to obtain a non-linear transformation from the native functional space to the standardized MNI space; regression of confound signals including the 6 motion parameters, motion outliers and average white matter and CSF signals. These average signals were obtained by registering a standardized mask to the native functional space by inverting the non-linear transformation obtained in the previous step. The residual images following nuisance regression were then used as signal of interest. These residuals were then registered to the standardized MNI space and temporally filtered between 0.01-0.08 Hz.

After pre-processing, regional time-series of BOLD signal were extracted using the Shen atlas (Soares et al., 2016) obtaining 268 time-series per-subject which were then used to crate matrices on Functional Connectivity (FC), by calculating the fisher transformation of the correlation of each pair of signals.

Analysis of FC was performed using the Network Based Statistics (NBS, https://www.nitrc.org/projects/nbs/, (Zalesky et al., 2010)) approach, performing a independent sample t-test to compare experimental and control groups. The NBS approach uses a GLM-style analysis and works in two steps: first, it regresses out the confounding factors and uses the residuals to test the hypothesis in each individual connection, obtaining the corresponding t-statistic; second, it estimates the significance of the obtained network. This is done by first calculating the size of the network of significant connections, thresholding it by a user defined value and counting the number of significant connections between interconnected nodes and comparing it to the size of networks obtained from 5000 random permutations of the groups. This procedure yields a non-parametric estimation of the significance of the network corrected for the Family-Wise Error rate. Because a user specified threshold determines the size of the network, and different thresholds can yield networks with different nature, a range of thresholds was tested (p = 0.01, 0.005, 0.001, 0.0005 and 0.0001), following the suggestion of the authors of the toolbox. Results were considered significant when the FWE corrected network p-value is ≤ 0.05. To help interpret possible significant networks we identified the key nodes by calculating the nodal t-value as the sum of all supra-threshold edges for each node.

Analysis of volumetric data was done using an independent sample t-test, controlling for sex, weight at birth and gestational age as well as total intra-cranial volume and corrected for multiple comparisons.

All comparisons were done controlling for sex, weight at birth and gestational age.

## 3. Results

### 3.1. Cohort characterization

The database is composed of 4607 parturitions, with 2243 females (48.7%) and 2364 males (51.2%), 4502 unifetal gestations (97.7%) and 105 gemelar gestations (2.3%). Mean gestational age at parturition was 38.7±1.7 weeks and 92.7% (4272) were term parturitions. Surprisingly, only in 2.4% (112) of the gestations, indication of prenatal sGC administration was present; this percentage is below the 8-9% estimated by the Instituto Nacional de Estatística (INE)/Statistics Portugal. Additional sGC-exposed infants were identified when evaluating the child clinical records in addition to maternal records. Of those indicated as prenatally sGC-exposed, 32.1% (36) were term gestations, 67% (75) were preterm and 1 had no reference to gestational age.

After assessing inclusion and exclusion criteria, 45 subjects of both sexes were enrolled in the study: 21 subjects of the sGC-exposed group (61.9% females) and 24 subjects of the non-exposed group (44% females). In the sGC-exposed group, mothers of 15 subjects were administered with two doses of intra-muscular 12mg of dexamethasone; 4 mothers had two doses of 6 mg of betamethasone and 2 mothers had 6 doses of 6mg of betamethasone. The reason for administration of sGC was risk of preterm labour (20 subjects) and placenta abruption (1 subject). Mean gestational age at sGCs administration was 32.4 weeks (28-34 weeks of gestation).

Individuals were selected to ensure that there was no major difference in the average gestational age (35.9 weeks in sGC-exposed group vs 36.7 weeks in non-exposed; Table 1). No differences were found in maternal age. sGC-exposed individuals presented reduced birth weight in comparison to non-exposed subjects (Table 1).

**Table 1.** Cohort details.

|  | **sGC-exposed**<br>**(n=21)** | **Non-exposed**<br>**(n=24)** | **Statistics**<br>**(p value)** |
| --- | --- | --- | --- |
| **Gestational Age (weeks)** | 35.8±2.2 | 36.6±2.7 | 0.235 |
| **Weight at birth (gr)** | 2419.4±632 | 3079±660 | **0.002\*** |
| **Current weight (kg)** | 58.9±9.7 | 63.4±10.1 | 0.243 |
| **Current height (m)** | 1.67±0.08 | 1.69±0.09 | 0.275 |
| **Body mass index** | 21.44±3.0 | 21.92±2.4 | 0.555 |
| **Ethnicity** | All Caucasian | All Caucasian | - |
| **Maternal Age at birth (years)** | 26.9±5.1 | 27.5±4.4 |  |
| **Paternal age at birth (years)** | 29.9±4.9 | 29.8±5.1 |  |
| **Maternal education (n)** |  |  | 0.683 |
| 4 years | 0 | 0 |  |
| 6 years | 4 | 3 |  |
| 9 years | 3 | 4 |  |
| 12 years | 6 | 6 |  |
| Bsc | 1 | 4 |  |
| *Licenciatura (4-5 year degree)* | 7 | 6 |  |
| Msc | 0 | 1 |  |
| PhD | 0 | 0 |  |
| **Paternal education (n)** |  |  | 0.921 |
| 4 years | 1 | 5 |  |
| 6 years | 7 | 7 |  |
| 9 years | 1 | 6 |  |
| 12 years | 5 | 4 |  |
| Bsc | 0 | 0 |  |
| *Licenciatura* | 6 | 2 |  |
| Msc | 0 | 0 |  |
| PhD | 1 | 0 |  |

### 3.2. Neuropsychological and psychiatric evaluation

BDI-II, BAI, PSS-10, SQD, SURPS, BIS/BAS average scores had no major significant differences between groups (Sup. Tables 1-5). There was evidence on subcomponent of the NEO-FFI test associating with sGC group, with sGC subjects being more prone to openness to experience than controls (Sup. Table 5).

### 3.3. Brain structural analysis in GC-exposed individuals

A subset of 19 sGC-exposed (64.7% female) and 11 non-exposed individuals (63.6% females) underwent the multimodal MRI at 17 years of age. There were no differences in gestational weeks at birth between the sGC-exposed (n=17) and non-exposed (n=11) groups (35.9±2.4 vs 36.5±3.3; p=0.579). Weight at birth was different between groups, with the sGC-exposed group presenting lower weight in comparison to the non-exposed group (2411.2±647 vs 3087±486 g; p=0.009). Results of the structural analysis are presented in Table 2. Here we present the p significance, t-value, mean value for each group and Hedge’s h value. Out of these results, none survive correction for multiple comparisons (a p<0.0019 for Family-Wise Error correction), but there is a robust trend for a bilateral enlargement in the caudate as well as the right superior frontal cortex of the sGC-exposed individuals (p=0.008 for both).

**Table 2.** Results from the comparison between sGC-exposed and non-exposed subjects on 12 bilateral volumes extracted from Freesurfer. Here we present the uncorrected p-value, the t-statistic value and hedge’s h value as a measure of effect size.

|  | p | t | h | p | t | h |
| --- | --- | --- | --- | --- | --- | --- |
| Region | left |  |  | right |  |  |
| Thalamus | 0.382 | 0.893 | 0.39 | 0.233 | 1.229 | 0.30 |
| Caudate | **0.008** | **-2.944** | **1.56** | **0.008** | **-2.932** | **1.53** |
| Putamen | 0.205 | -1.306 | 0.85 | 0.756 | -0.315 | 0.46 |
| Pallidum | 0.994 | 0.007 | 0.26 | 0.775 | 0.290 | 0.28 |
| Hippocampus | 0.127 | -1.591 | 1.01 | 0.143 | -1.522 | 0.83 |
| Amygdala | 0.708 | -0.380 | 0.26 | 0.189 | -1.356 | 0.27 |
| Accumbens | 0.915 | -0.108 | 0.08 | 0.682 | -0.416 | 0.29 |
| Anterior cingulum | 0.18 | -1.38 | 0.53 | 0.15 | -1.5 | 0.75 |
| Posterior cingulum | 0.18 | -1.37 | 0.72 | 0.11 | -1.7 | 0.81 |
| Superior Frontal | 0.015 | -2.66 | 1.12 | **0.008** | **-3.93** | **1.67** |
| Cerebellum | 0.81 | -0,24 | 0.34 | 0.81 | -0,25 | 0.41 |
| Precuneus | 0.278 | -1.113 | 0.65 | 0.215 | -1.280 | 0.84 |

### 3.4. Functional connectivity is altered in sGC-exposed adolescents

Comparing the sGC-exposed group to the non-exposed group using the NBS approach, and controlling for sex, gestational age at birth and weight birth, a network of altered functional connectivity (FC) was found with significant, or very nearly, p-values across all thresholds (Table 3, Figure 1). This network represented reduced FC in the sGC-exposed subjects (Figure 1D). At an edge threshold of *p* = 0.001, a network significance of *p* = 0.049 was found, over 66 connections, 58 nodes, with a large effect size *h* = 1.18. Within this network, the sGC-exposed group was found to have an average FC of 0.21 ± 0.10 and the control group 0.53 ± 0.18. This network involved primarily sub-cortical, cerebellar and frontal nodes. From the statistical centrality description of this network, we identified several nodes as being central to the altered FC network across the different thresholds including the Left (nodes #220 and #224) and Right (#84 and #88) Cingulate Gyrus, the Left ACC (#134) the Left Parahippocampal Gyrus (#231 and #233), the Right Precuneus (#42, #75 and #90), the Cerebellum (#71 and #247). The weight of all nodes involved in the affected networks across all thresholds can be found in Supplementary Table 6.

**Figure 1.**
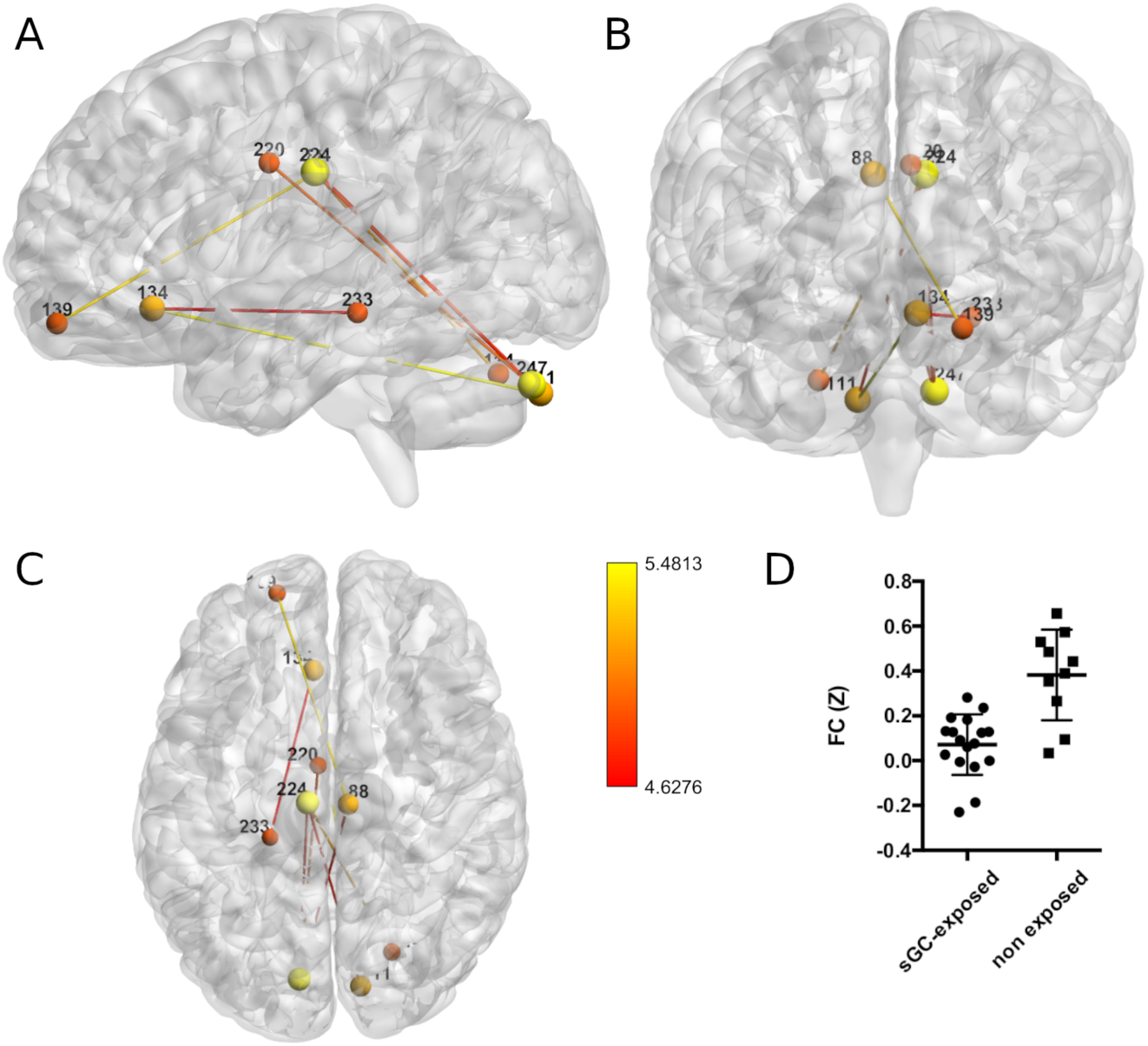
Network of reduced functional connectivity (FC) in sGC-exposed subjects, obtained using NBS at an edge threshold of p=0.001, with a network significance of p=0.049. Nodes and edges are color coded using a hot color scheme. We present sagittal (A), coronal (B) and axial (C) views of the brain and the (D) scatter plot of the average of FC for each group, bars represent mean and standard deviation values.

**Table 3.** Results from the NBS analysis at each edge threshold corrected for the FWE rate. Here are reported the network significance, number of edges involved and hedge’s h as a measure of effect size. Degrees of freedom = 25.

| Threshold (p/t-value) | P-network | N° edges | h |
| --- | --- | --- | --- |
| 0.01/2.79 | 0.041 | 699 | 0.96 |
| 0.005/3.08 | 0.041 | 361 | 1.03 |
| 0.001/3.73 | 0.049 | 66 | 1.18 |
| 0.0005/3.99 | 0.056 | 24 | 1.16 |
| 0.0001/4.62 | 0.026 | 8 | 1.32 |

## 4. Discussion

During gestation, the fetus is highly responsive to GCs, therefore maternal GC levels are tightly controlled. GCs are necessary for normative fetal brain development, however, exposure to excessive levels of these hormones may disrupt brain processes, leading to increased vulnerability to develop neuropsychiatric conditions later in life. Although a consensus is solid in that the administration of sGCs has decreased neonatal mortality and morbidity, there is still the need to perform additional studies to understand the long-term impact of prenatal sGC administration, as human studies are sparse.

In this study we investigated the long-term consequences of prenatal exposure to sGCs on the neuropsychological traits of adolescents, and evaluated its impact in the adolescent brain. We show that 17-year-old adolescents that were prenatally exposed to sGCs presented no major significant differences in the neuropsychological domains evaluated in this study. However, prenatal sGC-exposure was associated with changes in the functional connectivity of a resting state network, in particular in the Cingulate Gyrus, Left Parahippocampal Gyrus, the Right Precuneus, and the Cerebellum. Though no statistical differences were found in the volumes of these or other brain regions, as none of the results survive correction for multiple comparisons, a strong trend was observed for a bilateral enlargement in the caudate of the sGC-exposed individuals.

Several studies in animal models have shown that prenatal sGC exposure induces long-lasting molecular, structural, morphological, and functional alterations in different brain regions. For example, prenatal sGC exposure in rodents alters the HPA axis, leading to structural and functional changes in different brain regions, including the prefrontal cortex and limbic regions important for emotion processing and regulation such as the hippocampus, nucleus accumbens, and amygdala, among others (Cartier et al., 2016; McArthur et al., 2005; Oliveira et al., 2006, 2011, 2012; Reynolds, 2013; Rodrigues et al., 2012). Therefore, it is not surprising that animals prenatally exposed to sGC present emotional and behavioral deficits, including anhedonia and depressive-like behavior (Coimbra et al., 2017; Oliveira et al., 2006; Soares-Cunha et al., 2014), anxiety (Duarte et al., 2019; Oliveira et al., 2006) as well as increased addictive behaviors (Rodrigues et al., 2012).

In humans, studies are very limited, but some evidence suggests that prenatal sGC exposure is associated with emotional and behavioral changes later in life. For example, in a large cohort of >4500 mother-children dyads, the prevalence of mental, emotional and behavioral disturbances were higher in the sGC-exposed group when compared to non-exposed children (Wolford et al., 2019). A more recent study investigated a retrospective cohort of >670,000 children and revealed that sGC exposure was significantly associated with higher risk of any mental and behavioral disorder (Räikkönen et al., 2020). In our cohort we performed an extensive neuropsychological characterization using self-report questionnaires focusing on behavioral dimensions, emotional and personality traits. We found no significant differences between groups, with the exception of the openness to experience subcomponent of the NEO-FFI test, where the sGC-exposed group presented higher values. Regarding these results, several considerations have to be made. First, the small sample size may hinder any effects of sGC exposure. Second, we only used self-report questionnaires, in contrast with some of the previous studies. Third, individuals that had a family history of neuropsychiatric disorders, or had been diagnosed neuropsychiatric disorder, or that were treated for a psychiatric illness were excluded from the study, which could have biased the current results. However, also some other studies have found no association between prenatal sGC and later child outcomes, including hyperactivity, emotional symptoms, prosocial behavior, conduct, or peer problems (Stutchfield et al., 2013), or levels of affective problems or intelligence (Alexander et al., 2016; Davis et al., 2013). These contradictory findings can also result from different study designs, differences in measurements as well as differences in the regimen and timing of sGC administration.

Considering the importance of GCs for fetal brain development and maturation, it is not unexpected that prenatal exposure to sGC could lead to long-lasting brain alterations in humans. Surprisingly, to date, very few studies have evaluated brain structure, connectivity and/or functionality in sGC-exposed cohorts. An early brain imaging study showed that term-born infants exposed to multiple doses of prenatal sGC treatment exhibited reduced cortical folding (Modi et al., 2001). One study found that children (age 6-10 years) with a single course prenatal sGC exposure had prominent cortical thinning in the right anterior cingulate cortex (rACC), although they did not manifest significant affective problems at that age (REF). However, as a thinner rACC is associated with risk for affective problems, the authors postulated that this phenotype could arise later in life (Davis et al., 2013). Another study showed that the activity of the fronto-parietal network of sGC-exposed adolescents was reduced during conflict monitoring (Ilg et al., 2018). Herein, and after appropriate statistical correction, we did not find any significant structural changes in the brains of adolescents exposed to sGC during gestation. It is important to highlight that in our study the experimental group was contrasted with individuals that were not exposed to sCG during gestation but had a similar gestational age at birth.

In this study, and for the same experimental comparison, we showed that sGC-exposed individuals presented reduced FC of a network that involved primarily sub-cortical, cerebellar and frontal nodes, even after controlling for sex, gestational age at birth and weight birth. Several nodes were significantly altered, including the left ACC, which is in line with one of the few studies that exist that showed that prenatal sGC exposure led to prominent cortical thinning in the right ACC (Davis et al., 2013). In a task designed to evaluate cognitive conflict monitoring, a reduction was observed in the activation of the fronto-parietal network, which underlies cognitive and behavioral control, most notably in the cingulate cortex and precuneus in sGC-exposed adolescents (Ilg et al., 2018). Interestingly, both the ACC and precuneus functional connectivity were affected in our cohort, showing that these regions may be particularly affected by prenatal sGC exposure. In agreement, animal studies have shown that prenatal sGC exposure leads to structural changes in the ACC (Soares-Cunha et al., 2014).

Importantly, inevitably, our study also raises a clinically relevant question that regards the optimization of the current guidelines for sGC administration when a risk of preterm labour has been identified. A prospective cohort study showed that 20-60% of sGC treatments are prescribed to low risk of preterm labour women, of whom only 1–3% give birth within the next seven days (Wilms et al., 2015). In another study focusing in high-risk for preterm labour and consequent sGC administration, approximately one third to a half of pregnancies proceeded to term (Polyakov et al., 2007). This highlights the urgent need to identify reliable markers of preterm labour and indications for the use of sGC after a clear risk-benefit assessment.

### 4.1. Strengths and Limitations

A primary limitation of this study is the relatively small sample size which limits data interpretation and calls for additional studies to confirm the findings herein reported. Since birth weight is also a critical factor to consider, we also recognize that this cohort limited the comparison with individuals that were born full-term but displayed similar birth weights as the sGC-exposed group, and with individuals with similar gestational age and birth weight (that are particularly difficult to find). However, one relevant strength is the similarity in the gestational age at birth, with the inclusion of preterm individuals in the control group, which controls for prematurity-associated changes.

The other positive aspect is the comprehensive characterization of the cohort and the exclusion criteria, which limited the analysis to *healthy* individuals, so that additional factors such as previous health conditions and adverse events, that could contribute for the observed outcomes, could be *controlled* for.

Another limitation is the absence of maternal questionnaires during pregnancy to evaluate maternal depressive and anxious symptomatology, as these can influence maternal cortisol secretion and impact fetal development, as well as the absence of teacher and parental reports on adolescent behavior.

## 5. Conclusion

Even considering the study limitations, our results suggest that prenatal sGC exposure is associated with persistent functional, but not structural, changes in different brain regions, with no observable expression in the neuropsychological domains that we evaluated in adolescence. The clinical relevance of these findings remains to be determined. This study is of relevance, considering that ∼10% of pregnancies are at risk for preterm labour and sGC administration is the gold-standard treatment in these situations.

## 6. Disclosure summary

Authors have nothing to disclose. The funding bodies had no role in the design and conduct of the study; in the collection, analysis, and interpretation of the data; or in the preparation, review, or approval of the manuscript.

## 7. Funding

This work was funded by Foundation for Science and Technology (FCT) under the scope of the project PTDC/MED-NEU/29071/2017 (REWSTRESS) and POCI-01-0145-FEDER-016428 (MEDPERSYST). Part of the work was supported by a BIAL Foundation Grant (Bial Foundation 30/16).

## Supplementary Materials

**Supplementary Table 1:** Results of the Beck depression scale, Beck anxiety scale and Perceived Stress 10 scale. Data is presented as mean ± standard deviation (range).

| Scale | sGC-exposed group<br>M, SD (range) | Non-exposed group<br>M, SD (range) | Statistics |
| --- | --- | --- | --- |
| Beck depression inventory | 6.1±4.2 (1-17) | 4.9±3.7 (0-14) | p = .342<br>t(43) = 0.961 |
| Beck anxiety inventory | 8.8±6.4 (1-23) | 10.7±5.6 (1-21) | p = .282<br>t(43) = -1.09 |
| PSS-10 | 17.8±3.3 (11-23) | 18.2±2.5 (14-24) | p = .691<br>t(43) = -0.401 |

**Supplementary Table 2:** Results of the Strengths and Difficulties Questionnaire (SDQ). Data is presented as mean ± standard deviation (range).

| SDQ Component | sGC-exposed group<br>M, SD (range) | Non-exposed group<br>M, SD (range) | Statistics |
| --- | --- | --- | --- |
| Emotional symptoms | 3.3±2.4 (0-7) | 3.8±2.1 (1-9) | p = .502<br>t(43) = -0.677 |
| Hyperactivity | 3.5±1.5 (1-7) | 3.9±2 (1-8) | p = .518<br>t(43) = -0.652 |
| Conduct problems | 1.4±0.9 (0-3) | 1.5±1.4 (0-5) | p = .834<br>t(43) = -0.211 |
| Peer problems | 3.1±0.9 (2-5) | 2.8±1 (1-5) | p = .396<br>t(43) = 0.857 |
| Pro-social | 8.1 (2-10) | 9.1 (6-10) | p = .71<br>U = 177<br>Z = -1.085 |
| Total Score | 11.3±3.4 (6-17) | 11.9±4.4 (5-22) | p = .593<br>t(43) = -0.539 |

**Supplementary Table 3:** Results of the Substance Use Risk Profile Scale. Data is presented as mean ± standard deviation (range).

| Personality type | sGC-exposed group<br>M, SD (range) | Non-exposed group<br>M, SD (range) | Statistics |
| --- | --- | --- | --- |
| Hopelessness | 21.5±2.5 (16-25) | 21.5±2.1 (17-25) | p = .973<br>t(43) = 0.034 |
| Anxiety sensation | 11.2±3.2 (5-18) | 10.8±2.3 (6-15) | p = .665<br>t(43) = 0.436 |
| Sensation seeking | 17.1±3.2 (10-21) | 17.8±3 (12-23) | p = .402<br>t(43) = -0.847 |
| Impulsivity | 10.3±2.5 (5-15) | 10.8±2.5 (7-17) | p = .468<br>t(43) = -0.732 |

**Supplementary Table 4:** Results of the Behavioral Inhibition Scale/Behavioral Avoidance Scale (BIS/BAS). Data is presented as mean ± standard deviation (range).

| BIAS/BAS Component | sGC-exposed group<br>M, SD (range) | Non-exposed group<br>M, SD (range) | Statistics |
| --- | --- | --- | --- |
| BIS | 17.8±5.4 (7-27) | 17.2±4.2 (9-25) | p = .654<br>t(43) = 0.451 |
| BAS Drive | 10.5±1.6 (8-14) | 10.2±2.6 (5-15) | p = .688<br>t(43) = 0.404 |
| BAS Fun Seeking | 10.2±2.5 (4-14) | 10±3 (6-16) | p = .866<br>t(43) = 0.17 |
| BAS Reward responsiveness | 12.3±5.6 (5-20) | 11.3±3.6 (5-20) | p = .6<br>t(43) = 0.529 |

**Supplementary Table 5:** Results of the NEO-FFI. Data is presented as mean ± standard deviation (range).

| SDQ Component | sGC-exposed group<br>M, SD (range) | Non-exposed group<br>M, SD (range) | Statistics |
| --- | --- | --- | --- |
| Extraversion | 9.8 ±2.7 (5-16) | 10.8±2.4 (6-14) | p = .178<br>t(43) = -1.371 |
| Agreeableness | 9.2±2.4 (5-14) | 9±2.8 (4-16) | p = .771<br>t(43) = 0.293 |
| Consciousness | 7.4±2.2 (3-11) | 7.5±2.1 (3-12) | p = .904<br>t(43) = -0.121 |
| Neuroticism | 5.6±2.5 (0-12) | 5.4±2 (2-11) | p = .765<br>t(43) = 0.3 |
| Openness to experience | 9.7±3.1(3-16) | 7.4±3 (2-13) | **p = .015\***<br>t(43) = 2.54 |

**Supplementary Table 6:** Sum of t-value weight of each node for the threshold-surviving network for all tested edge thresholds. Here we report the sum of the t-statistic value over all significant edges of each node, with higher values representing a stronger involvement of the node within the affected network. The top of the columns for networks that survived corrected significance threshold are highlighted in bold. Empty cells represent sum values of 0.

| # | ROI | region | threshold p value |  |  |  |  |
| --- | --- | --- | --- | --- | --- | --- | --- |
|  |  |  | **0.01** | **0.005** | **0.001** | **0.0005** | **0.0001** |
| 1 | Right | Superior Frontal Gyrus | 6.79 | 3.73 | 3.73 |  |  |
| 2 | Right | Rectal Gyrus | 6.53 | 6.53 |  |  |  |
| 3 | Right | Medial Frontal Gyrus | 11.93 | 3.28 |  |  |  |
| 4 | Right | Inferior Frontal Gyrus | 4.14 | 4.14 | 4.14 | 4.14 |  |
| 5 | Right | Anterior Cingulate | 12.88 | 6.92 | 3.80 |  |  |
| 6 | Right | Medial Frontal Gyrus | 6.92 | 6.92 |  |  |  |
| 7 | Right | Superior Frontal Gyrus | 5.96 | 3.11 |  |  |  |
| 8 | Right | Middle Frontal Gyrus | 49.71 | 32.35 | 8.51 |  |  |
| 9 | Right | Middle Frontal Gyrus | 12.93 | 12.93 |  |  |  |
| 10 | Right | Superior Frontal Gyrus | 16.27 | 10.31 |  |  |  |
| 11 | Right | Superior Frontal Gyrus | 30.90 | 13.57 | 4.05 |  |  |
| 13 | Right | Middle Frontal Gyrus | 24.68 | 7.07 | 3.94 |  |  |
| 14 | Right | Middle Frontal Gyrus | 9.46 | 3.75 | 3.75 |  |  |
| 15 | Right | Cingulate Gyrus | 21.83 | 13.30 |  |  |  |
| 16 | Right | Inferior Frontal Gyrus | 5.87 |  |  |  |  |
| 17 | Right | Middle Frontal Gyrus | 3.72 | 3.72 |  |  |  |
| 18 | Right | Inferior Frontal Gyrus | 45.39 | 13.15 |  |  |  |
| 19 | Right | Middle Frontal Gyrus | 42.67 | 19.14 |  |  |  |
| 21 | Right | Inferior Frontal Gyrus | 12.14 | 6.35 |  |  |  |
| 22 | Right | Middle Frontal Gyrus | 3.61 | 3.61 |  |  |  |
| 24 | Right | Medial Frontal Gyrus | 2.85 |  |  |  |  |
| 25 | Right | Medial Frontal Gyrus | 24.85 | 19.21 |  |  |  |
| 27 | Right | Precentral Gyrus | 10.03 | 7.00 |  |  |  |
| 29 | Right | Superior Frontal Gyrus | 2.97 |  |  |  |  |
| 30 | Right | Superior Frontal Gyrus | 37.60 | 23.51 |  |  |  |
| 31 | Right | Precentral Gyrus | 20.54 | 3.12 |  |  |  |
| 32 | Right | Precentral Gyrus | 15.92 | 10.14 |  |  |  |
| 33 | Right | Postcentral Gyrus | 15.19 | 9.49 |  |  |  |
| 34 | Right | Extra-Nuclear<br>Broadman area<br>13 | 2.84 |  |  |  |  |
| 35 | Right | Insula | 10.14 | 7.22 |  |  |  |
| 36 | Right | Inferior Frontal<br>Gyrus | 3.22 | 3.22 |  |  |  |
| 38 | Right | Inferior Parietal<br>Lobule | 60.70 | 34.83 | 8.62 |  |  |
| 39 | Right | Postcentral<br>Gyrus | 18.02 | 3.52 |  |  |  |
| 40 | Right | Insula | 9.00 | 3.20 |  |  |  |
| 41 | Right | Postcentral<br>Gyrus | 87.75 | 54.93 | 12.01 |  |  |
| 42 | Right | Precuneus | 63.66 | 31.86 | 12.18 | 8.22 |  |
| 43 | Right | Superior Parietal<br>Lobule | 6.33 | 3.50 |  |  |  |
| 44 | Right | Precuneus | 6.54 | 3.62 |  |  |  |
| 45 | Right | Postcentral<br>Gyrus | 7.37 | 7.37 | 3.83 |  |  |
| 46 | Right | Postcentral<br>Gyrus | 6.13 | 3.18 |  |  |  |
| 47 | Right | Inferior Parietal<br>Lobule | 6.96 | 6.96 |  |  |  |
| 48 | Right | Angular Gyrus | 6.28 | 3.35 |  |  |  |
| 49 | Right | Superior<br>Occipital Gyrus | 3.07 |  |  |  |  |
| 50 | Right | Superior<br>Temporal Gyrus | 17.34 | 14.31 | 4.24 | 4.24 |  |
| 51 | Right | Superior<br>Temporal Gyrus | 9.21 | 3.56 |  |  |  |
| 52 | Right | Superior<br>Temporal Gyrus | 27.71 | 13.16 |  |  |  |
| 53 | Right | Superior<br>Temporal Gyrus | 12.13 | 3.37 |  |  |  |
| 55 | Right | Inferior Temporal<br>Gyrus | 9.18 | 6.28 |  |  |  |
| 56 | Right | Inferior Temporal<br>Gyrus | 19.45 | 13.59 | 4.17 | 4.17 |  |
| 58 | Right | Inferior Temporal<br>Gyrus | 11.85 | 3.09 |  |  |  |
| 59 | Right | Fusiform Gyrus | 3.16 | 3.16 |  |  |  |
| 61 | Right | Superior<br>Temporal Gyrus | 24.93 | 13.10 |  |  |  |
| 62 | Right | Parietal<br>Operculum | 3.08 | 3.08 |  |  |  |
| 63 | Right | Superior Temporal Gyrus | 6.45 | 6.45 |  |  |  |
| 64 | Right | Inferior Temporal Gyrus | 16.79 | 13.77 |  |  |  |
| 65 | Right | Superior Temporal Gyrus | 2.82 |  |  |  |  |
| 66 | Right | Fusiform Gyrus | 33.70 | 21.90 | 8.37 |  |  |
| 67 | Right | Declive of the Cerebellum | 6.50 | 6.50 |  |  |  |
| 68 | Right | Culmen of the Cerebellum | 3.00 |  |  |  |  |
| 69 | Right | Inferior Temporal Gyrus | 19.92 | 10.93 | 7.74 |  |  |
| 70 | Right | Inferior Temporal Gyrus | 46.86 | 20.55 | 3.93 |  |  |
| 71 | Right | Culmen of the Cerebellum | 59.50 | 42.17 | 12.11 |  |  |
| 72 | Right | Declive of the Cerebellum | 2.86 |  |  |  |  |
| 73 | Right | Middle Occipital Gyrus | 6.57 | 3.67 |  |  |  |
| 74 | Right | Inferior Occipital Gyrus | 14.84 |  |  |  |  |
| 75 | Right | Precuneus | 42.89 | 39.84 | 23.84 |  |  |
| 76 | Right | Lingual Gyrus Gray | 16.17 | 10.42 |  |  |  |
| 77 | Right | Cuneus Gray | 5.90 |  |  |  |  |
| 78 | Right | Middle Occipital Gyrus | 3.20 | 3.20 |  |  |  |
| 81 | Right | Inferior Occipital Gyrus | 103.39 | 59.51 |  |  |  |
| 82 | Right | Cuneus | 8.74 |  |  |  |  |
| 83 | Right | Anterior Cingulate | 12.24 | 6.48 |  |  |  |
| 84 | Right | Cingulate Gyrus | 36.71 | 25.04 | 8.30 |  |  |
| 85 | Right | Posterior Cingulate | 28.29 | 10.71 | 4.35 | 4.35 |  |
| 86 | Right | Posterior Cingulate | 29.97 | 9.80 |  |  |  |
| 87 | Right | Parahippocampal Gyrus | 6.04 |  |  |  |  |
| 88 | Right | Cingulate Gyrus | 156.30 | 121.56 | 54.74 | 32.02 | 10.09 |
| 89 | Right | Paracentral Lobule | 21.53 | 7.01 |  |  |  |
| 90 | Right | Precuneus | 90.08 | 57.82 | 8.12 | 8.12 |  |
| 91 | Right | Paracentral Lobule | 9.38 | 3.54 |  |  |  |
| 94 | Right | Parahippocampal Gyrus | 2.82 |  |  |  |  |
| 95 | Right | Parahippocampal Gyrus | 6.59 | 3.59 |  |  |  |
| 96 | Right | Parahippocampal Gyrus | 19.11 | 7.42 | 4.30 |  |  |
| 98 | Right | Parahippocampal Gyrus | 42.06 | 13.20 |  |  |  |
| 99 | Right | Parahippocampal Gyrus | 16.16 | 13.20 |  |  |  |
| 100 | Right | Inferior Semi-Lunar Lobule | 10.42 | 7.63 | 3.91 |  |  |
| 101 | Right | Culmen of the Cerebellum | 6.15 | 3.29 |  |  |  |
| 102 | Right | Crus I | 19.65 | 7.95 | 4.31 | 4.31 |  |
| 104 | Right | Cerebellar Tonsil | 6.38 | 6.38 |  |  |  |
| 106 | Right | Declive of the Cerebellum | 2.90 |  |  |  |  |
| 107 | Right | Cerebellar Tonsil | 14.18 | 14.18 | 7.78 |  |  |
| 108 | Right | Cerebellar Area | 25.26 | 16.25 |  |  |  |
| 111 | Right | Cerebellar Pyramis | 31.24 | 25.45 | 18.65 | 18.65 | 10.21 |
| 112 | Right | Cerebellar Area | 25.18 | 7.29 |  |  |  |
| 113 | Right | Cerebellar Tonsil | 9.29 | 3.44 |  |  |  |
| 114 | Right | Cerebellar Pyramis | 11.84 | 11.84 | 5.22 | 5.22 | 5.22 |
| 115 | Right | Cerebellar Area | 3.35 | 3.35 |  |  |  |
| 116 | Right | Cerebellar Area | 11.62 | 3.09 |  |  |  |
| 117 | Right | Cerebellar Tonsil | 2.97 |  |  |  |  |
| 121 | Right | Caudate Body | 3.51 | 3.51 |  |  |  |
| 126 | Right | Thalamus | 35.42 | 20.86 | 7.99 |  |  |
| 128 | Right | Thalamus | 6.00 |  |  |  |  |
| 129 | Right | Cerebellar Area | 5.99 |  |  |  |  |
| 130 | Right | Upper Brainstem Area | 9.30 | 6.41 |  |  |  |
| 132 | Right | Upper Brainstem Area | 2.89 |  |  |  |  |
| 133 | Right | Cerebellar Tonsil | 2.83 |  |  |  |  |
| 134 | Left | Anterior Cingulate | 35.58 | 23.96 | 10.11 | 10.11 | 10.11 |
| 135 | Left | Inferior Frontal Gyrus | 3.63 | 3.63 |  |  |  |
| 136 | Left | Rectal Gyrus | 5.88 |  |  |  |  |
| 137 | Left | Rectal Gyrus | 15.54 | 4.03 |  |  |  |
| 138 | Left | Medial Frontal Gyrus | 6.22 | 3.35 |  |  |  |
| 139 | Left | Superior Frontal Gyrus | 33.94 | 16.40 | 9.95 | 9.95 | 5.37 |
| 140 | Left | Medial Frontal Gyrus | 8.74 |  |  |  |  |
| 141 | Left | Medial Frontal Gyrus | 20.27 | 14.22 | 3.82 |  |  |
| 142 | Left | Middle Frontal Gyrus | 51.27 | 24.98 | 11.71 |  |  |
| 143 | Left | Middle Frontal Gyrus | 18.90 | 13.02 |  |  |  |
| 144 | Left | Middle Frontal Gyrus | 11.50 |  |  |  |  |
| 145 | Left | Superior Frontal Gyrus | 15.77 | 6.64 |  |  |  |
| 146 | Left | Middle Frontal Gyrus | 15.03 | 3.45 |  |  |  |
| 147 | Left | Middle Frontal Gyrus | 19.56 | 10.90 | 4.16 |  |  |
| 149 | Left | Superior Frontal Gyrus | 6.30 | 3.28 |  |  |  |
| 150 | Left | Medial Frontal Gyrus | 29.67 | 20.85 |  |  |  |
| 151 | Left | Inferior Frontal Gyrus | 3.07 |  |  |  |  |
| 152 | Left | Middle Frontal Gyrus | 12.29 | 3.47 |  |  |  |
| 153 | Left | Inferior Frontal Gyrus | 6.24 | 3.44 |  |  |  |
| 154 | Left | Middle Frontal Gyrus | 36.54 | 25.10 | 12.11 | 8.28 |  |
| 155 | Left | Insula | 17.02 | 14.18 |  |  |  |
| 156 | Left | Inferior Frontal Gyrus | 34.49 | 31.65 | 9.03 |  |  |
| 157 | Left | Inferior Frontal Gyrus | 22.78 | 10.94 | 4.03 |  |  |
| 158 | Left | Precentral Gyrus | 12.02 |  |  |  |  |
| 159 | Left | Precentral Gyrus | 5.68 |  |  |  |  |
| 160 | Left | Precentral Gyrus | 19.32 | 10.58 | 3.75 |  |  |
| 161 | Left | Cingulate Gyrus | 2.89 |  |  |  |  |
| 162 | Left | Superior Frontal Gyrus | 41.26 | 20.83 |  |  |  |
| 163 | Left | Precentral Gyrus | 10.58 | 10.58 |  |  |  |
| 164 | Left | Superior Frontal Gyrus | 2.87 |  |  |  |  |
| 165 | Left | Precentral Gyrus | 28.39 | 17.04 |  |  |  |
| 166 | Left | Middle Frontal Gyrus | 15.35 | 9.70 |  |  |  |
| 167 | Left | Precentral Gyrus | 5.88 |  |  |  |  |
| 168 | Left | Insula | 31.07 | 10.66 |  |  |  |
| 169 | Left | Insula | 2.89 |  |  |  |  |
| 170 | Left | Insula-Clastrum | 9.24 | 3.41 |  |  |  |
| 171 | Left | Postcentral Gyrus | 9.01 | 3.11 |  |  |  |
| 172 | Left | Postcentral Gyrus | 12.25 | 3.47 |  |  |  |
| 173 | Left | Insula | 9.40 | 3.75 |  |  |  |
| 175 | Left | Superior Parietal Lobule | 44.48 | 23.99 | 3.98 |  |  |
| 176 | Left | Cuneus | 40.49 | 20.27 |  |  |  |
| 177 | Left | Precuneus | 3.11 | 3.11 |  |  |  |
| 178 | Left | Superior Parietal Lobule | 56.88 | 51.21 | 17.57 |  |  |
| 179 | Left | Inferior Parietal Lobule | 18.35 | 6.81 |  |  |  |
| 180 | Left | Superior Temporal Gyrus | 34.14 | 19.32 |  |  |  |
| 181 | Left | Postcentral Gyrus | 12.18 | 6.34 |  |  |  |
| 182 | Left | Inferior Parietal Lobule | 8.83 |  |  |  |  |
| 183 | Left | Superior Temporal Gyrus | 8.92 | 3.15 |  |  |  |
| 184 | Left | Inferior Parietal Lobule | 9.20 | 3.46 |  |  |  |
| 185 | Left | Middle Temporal Gyrus | 9.56 | 6.60 |  |  |  |
| 186 | Left | Superior Temporal Gyrus | 78.75 | 46.57 |  |  |  |
| 189 | Left | Superior Temporal Gyrus | 2.96 |  |  |  |  |
| 190 | Left | Fusiform Gyrus | 2.89 |  |  |  |  |
| 191 | Left | Superior Temporal Gyrus | 9.68 | 6.85 |  |  |  |
| 192 | Left | Middle Temporal Gyrus | 17.49 | 14.69 |  |  |  |
| 193 | Left | Inferior Temporal Gyrus | 12.05 | 3.41 |  |  |  |
| 194 | Left | Inferior Temporal Gyrus | 9.18 | 3.36 |  |  |  |
| 195 | Left | Uncus | 21.13 | 9.55 |  |  |  |
| 196 | Left | Inferior Temporal Gyrus | 14.65 | 3.13 |  |  |  |
| 197 | Left | Middle Temporal Gyrus | 36.07 | 24.31 |  |  |  |
| 198 | Left | Culmen of the Cerebellum | 15.74 | 7.01 |  |  |  |
| 199 | Left | Inferior Temporal Gyrus | 89.14 | 51.04 | 21.13 |  |  |
| 200 | Left | Fusiform Gyrus | 31.26 | 11.02 | 3.90 |  |  |
| 201 | Left | Culmen of the Cerebellum | 61.96 | 38.32 | 7.72 |  |  |
| 203 | Left | Middle Temporal Gyrus | 44.83 | 24.76 | 8.23 | 8.23 |  |
| 206 | Left | Fusiform Gyrus | 11.88 |  |  |  |  |
| 207 | Left | Declive of the Cerebellum | 20.10 | 11.47 |  |  |  |
| 208 | Left | Cuneus | 27.72 | 15.97 |  |  |  |
| 209 | Left | Inferior Temporal Gyrus | 12.26 | 6.35 |  |  |  |
| 210 | Left | Inferior Occipital Gyrus | 6.17 | 3.35 |  |  |  |
| 211 | Left | Lingual Gyrus | 50.57 | 15.35 |  |  |  |
| 212 | Left | Cuneus | 19.47 | 13.58 |  |  |  |
| 214 | Left | Inferior Occipital Gyrus | 5.90 |  |  |  |  |
| 215 | Left | Cuneus | 6.31 |  |  |  |  |
| 216 | Left | Posterior Cingulate | 8.72 |  |  |  |  |
| 218 | Left | Medial Frontal Gyrus | 22.32 | 16.45 |  |  |  |
| 219 | Left | Cingulate Gyrus | 5.70 |  |  |  |  |
| 220 | Left | Cingulate Gyrus | 118.91 | 60.73 | 17.29 | 17.29 | 4.94 |
| 221 | Left | Cingulate Gyrus | 32.77 | 18.14 | 4.91 |  |  |
| 222 | Left | Posterior Cingulate | 25.33 | 16.73 |  |  |  |
| 223 | Left | Cingulate Gyrus | 20.97 | 9.69 |  |  |  |
| 224 | Left | Cingulate Gyrus | 85.30 | 67.70 | 27.02 | 19.31 | 14.73 |
| 225 | Left | Precuneus | 15.04 | 3.31 |  |  |  |
| 226 | Left | Precuneus | 19.39 | 10.47 | 4.00 |  |  |
| 227 | Left | Posterior Cingulate | 30.67 | 13.37 |  |  |  |
| 228 | Left | Parahippocampal Gyrus | 22.74 | 16.80 |  |  |  |
| 231 | Left | Parahippocampal Gyrus | 149.21 | 108.23 | 20.22 |  |  |
| 233 | Left | Parahippocampal Gyrus | 56.75 | 33.44 | 12.84 | 8.97 | 4.63 |
| 234 | Left | Parahippocampal Gyrus | 23.28 | 6.21 |  |  |  |
| 236 | Left | Inferior Semi-Lunar Lobule | 9.38 | 3.42 |  |  |  |
| 237 | Left | Cerebellar Tonsil | 2.82 |  |  |  |  |
| 238 | Left | Cerebellar Tonsil | 60.39 | 31.20 | 8.40 |  |  |
| 239 | Left | Cerebellar Area | 16.66 | 13.63 |  |  |  |
| 241 | Left | Crus I | 3.14 | 3.14 |  |  |  |
| 242 | Left | Inferior Semi-Lunar Lobule | 4.26 | 4.26 | 4.26 | 4.26 |  |
| 243 | Left | Cerebellar Area | 9.61 | 9.61 |  |  |  |
| 244 | Left | Culmen of the Cerebellum | 2.89 |  |  |  |  |
| 245 | Left | Culmen of the Cerebellum | 14.85 | 3.12 |  |  |  |
| 246 | Left | Cerebellar Tonsil | 9.22 |  |  |  |  |
| 247 | Left | Uvula of the Cerebellum | 17.40 | 14.44 | 14.44 | 14.44 | 14.44 |
| 248 | Left | Declive of the Cerebellum | 20.51 | 14.65 | 7.78 |  |  |
| 251 | Left | Upper Brainstem Area | 3.52 | 3.52 |  |  |  |
| 252 | Left | Cerebellar Tonsil | 12.28 | 3.65 |  |  |  |
| 253 | Left | Pyramis of the Cerebellum | 8.89 | 3.15 |  |  |  |
| 254 | Left | Cerebellar Area | 2.84 |  |  |  |  |
| 255 | Left | Cerebellar Area | 19.27 | 16.21 |  |  |  |
| 258 | Left | Caudate | 12.06 | 12.06 | 8.42 | 4.53 |  |
| 259 | Left | Accumbens | 12.07 | 6.36 |  |  |  |
| 262 | Left | Thalamus | 33.89 | 25.23 | 11.63 |  |  |
| 266 | Left | Brain Stem Area | 3.31 |  |  |  |  |
| 267 | Left | Brain Stem Area | 31.90 | 23.02 | 16.47 | 12.57 |  |
| 268 | Left | Brain Stem Area | 27.76 | 24.81 | 4.24 | 4.24 |  |

## Notes

### Competing Interest Statement

The authors have declared no competing interest.

